# Differential Impact of Biomechanical Constraints on Control Signal Dimensionality for Gravity Support Versus Propulsion

**DOI:** 10.1101/2024.02.12.579990

**Authors:** AS Korol, V Gritsenko

## Abstract

Neural control of movement has to overcome the problem of redundancy in the multidimensional musculoskeletal system. The problem can be solved by reducing the dimensionality of the control space of motor commands, i.e., through muscle synergies or motor primitives. Evidence for this solution exists, multiple studies have obtained muscle synergies using decomposition methods. These synergies vary across different workspaces and are present in both dominant and non-dominant limbs. Here we explore the dimensionality of control space by examining muscle activity patterns across reaching movements in different directions starting from different postures performed bilaterally by healthy individuals. We further explore the effect of biomechanical constraints on the dimensionality of control space. We are building on top of prior work showing that muscle activity profiles can be explained by applied moments about the limb joints that reflect the biomechanical constraints. These muscle torques derived from motion capture represent the combined actions of muscle contractions that are under the control of the nervous system. Here we test the generalizability of the relationship between muscle torques and muscle activity profiles across different starting positions and between limbs. We also test a hypothesis that the dimensionality of control space is shaped by biomechanical constraints. We used principal component analysis to evaluate the contribution of individual muscles to producing muscle torques across different workspaces and in both dominant and non-dominant limbs. Results generalize and support the hypothesis. We show that the muscle torques that support the limb against gravity are produced by more consistent combinations of muscle co-contraction than those that produce propulsion. This effect was the strongest in the non-dominant arm moving in the lateral workspace on one side of the body.

## Introduction

The production of movement involves integrating biomechanical, neural, and environmental factors. Muscle recruitment by the central nervous system (CNS) causes motion or lack thereof, such as when maintaining posture, through the nonlinear production of forces by muscles about the degrees of freedom (DOFs) of the joints. Body inertia, the number of DOFs, and external forces then shape the resulting posture and movement. This biomechanics is complex enough that neural sensorimotor circuits must embed its dynamics for efficient and robust control [1–3]. However, a problem of redundancy exists, i.e., the problem of choosing among multiple muscles and combinations of joint angles that are possible for a given desired hand position or motion. This problem may be resolved by reducing the dimensionality of the space of motor commands by the CNS, a concept known as muscle synergies or motor primitives. This concept implies that muscles (or motoneurons in the spinal cord) are recruited in groups that represent a specific action so that a combination of the smaller number of synergies can produce a larger number of different movements [4–8]. The anatomical organization of muscles that form agonistic and antagonistic groups that can be activated by fewer control signals constraining the dimensionality of control space [9]. The spatial organization of motoneuron pools in the spinal cord also captures the functional relationships of the muscles they innervate, further supporting the idea of reduced dimensionality of control space [10]. Indeed, the neural activation of muscles observed with surface electromyography (EMG) is reducible to a low dimensional space for certain types of movements, including reaching [4,8,11]. However, the dimensionality reduction can be done using different mathematical methods applied to data organized in different ways [12,13]. These methods all obtain control space solutions with lower dimensions than the original dataset, but their neuromuscular or biomechanical underpinnings are not always clear.

Animal studies have worked out the basic modular and hierarchal organization of the neural motor control system that comprises nested feedback loops from sensory afferents to spinal motor neurons that encompass progressively more delayed processing circuits including the spinal reflexes, reticular formation in the brainstem, and primary sensory and motor cortices in the brain to name a few [14,15] (Fig 1A). Relating the synergies obtained with decomposition methods to the dynamic neural control units is not trivial. The commonly used Principal Component Analysis (PCA) can be applied across muscles, task conditions, and time in two ways, to obtain temporally-variable synergies (temporal synergies) or temporally invariant synergies (spatial synergies) [16]. A recent study has shown that the temporal dynamics of the spinal central pattern generator that controls rhythmic movements is captured well using temporal synergies obtained with PCA of EMG during locomotion [17]. We have also shown that the temporal dynamics of muscle torques needed to support the limb against gravity and for propulsion are well described by the 1^st^ and 2^nd^ temporal synergies respectively obtained with PCA. These two components of muscle action may rely on postural mechanisms governed by the pontomedullary reticular formation and the phasic velocity-dependent output of the primary motor cortex respectively [18–21]. For reaching, the temporal synergies have been shown to be more accurately capturing the temporal aspects of muscle activity and potentially neural control dimensionality [16].

**Figure 1.**
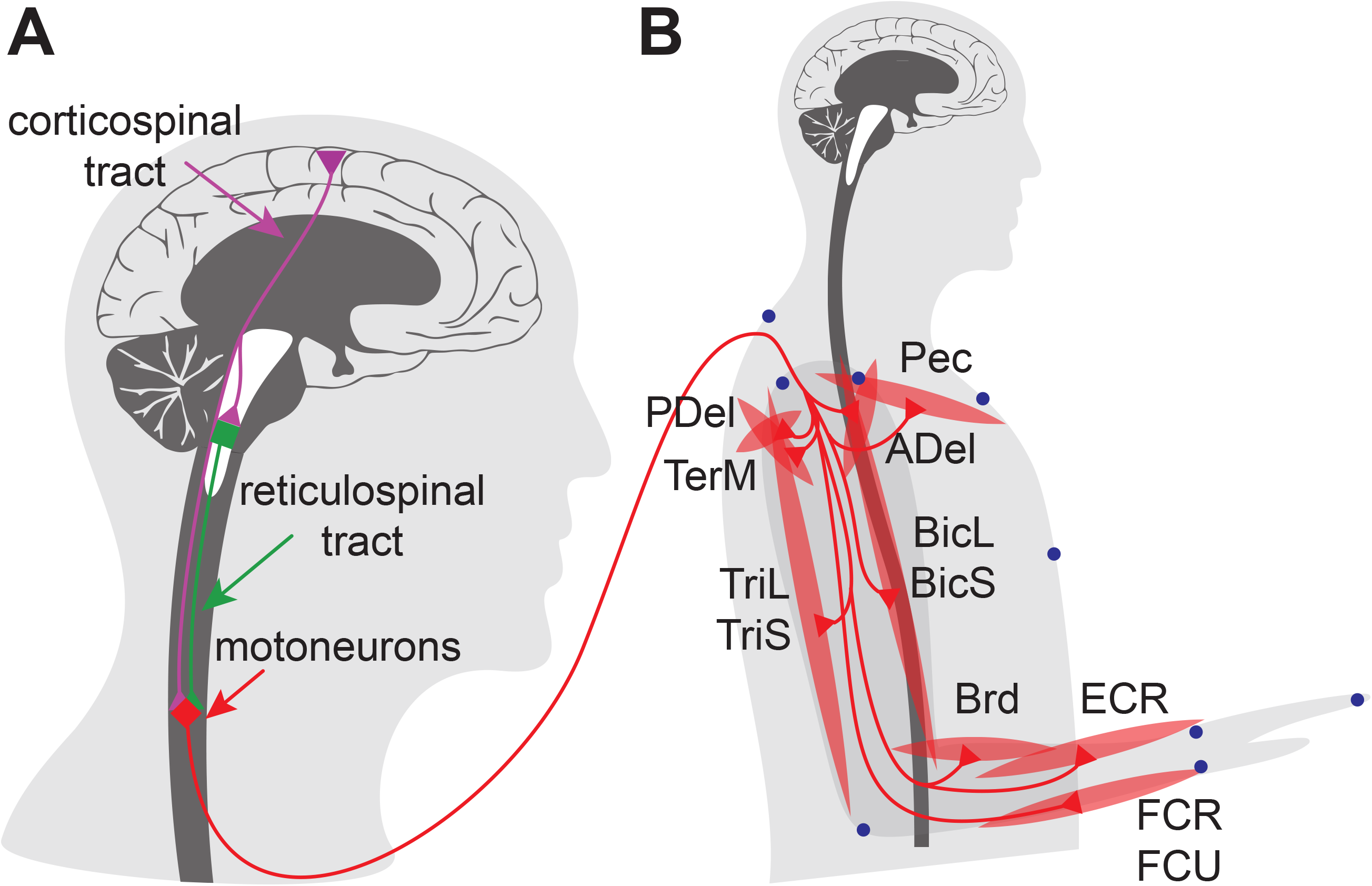
Schematic of neuromuscular system. **A.** Basic components of the neural circuit involved in muscle activation. Drawing is not to scale. **B.** Innervation of muscles recorded during the experiment and their arrangement on the arm. Abbreviations are as follows: the clavicular head of pectoralis (Pec), teres major (TerM), anterior deltoid (ADel), posterior deltoid (PDel), the long and lateral heads of triceps (TriL and TriS), the short and long heads of biceps (BiS and BiL), brachioradialis (Brd), flexor capri radialis (FCR), flexor carpi ulnaris (FCU), and extensor capri radialis (ECR). Blue dots indicate locations of LED markers for motion capture.

The temporal profile of the envelope of surface EMG is closely related to the force the corresponding muscle is producing in response to neural activation [22]. More recently we have shown that for reaches with the dominant hand towards visual targets in three-dimensional space, the EMG profiles are closely related to the muscle torques that cause motion [23]. Specifically, the static component of EMG that underlies postural forces needed to support the arm in specific position and during transitions between positions is closely related to the components of muscle torques that include gravity terms in the equations of motion. This suggests that the static component of EMG envelope from a given muscle reflects the muscle’s contribution to counteracting gravity load on the joints it spans. Moreover, the residual phasic component of EMG is closely related to the residual muscle torque that underlies the acceleration and deceleration forces toward the reaching goal after gravity-related component is subtracted. This suggests that the phasic component of EMG envelope from a given muscle reflects the muscle’s contribution to propulsion toward the target and stopping there. Here we will test the generalizability of these relationships between muscle activity and torque profiles to the non-dominant limb and across different workspaces.

Dimensionality reduction of control space can be observed through the co-contraction of multiple muscles. For example, a single control signal can activate both the brachioradialis and biceps muscles to produce an elbow flexion moment. This is the co-contraction of agonistic muscles. Moreover, that same control signal activating the triceps muscle, which opposes the action of the elbow flexors mentioned above, will both reduce the elbow flexion torque and increase the stiffness of elbow and shoulder joints. This is the co-contraction of antagonistic muscles. Both types of co-contraction are observed during reaching. The agonistic co-contraction together with reciprocal activation of antagonistic muscles contribute directly to muscle torques computed with inverse models and are likely facilitated through spinal reflexes and CPG. For example, stimulation of the primary afferents from the medial gastrocnemius muscle produces EPSPs not only in the motoneurons innervating that same muscle but also in motoneurons innervating lateral gastrocnemius and soleus, contributing to agonistic co-contraction [24]. The levels of antagonistic co-contraction are thought to vary with the dynamical requirements of the movement that vary with reaching in different directions from different starting postures [21,25]. While the CNS is thought to control whole limb stiffness through co-contraction [26,27], we do not know how this co-contraction signal is generated by the nervous system to help reduce the dimensionality of control space.

In this study, healthy human participants performed unconstrained pointing movements toward visual targets located equidistantly from one of two central targets. The reaching movements in multiple directions were performed withing the typical reaching workspace by either dominant (right) or non-dominant (left) arm. We collected motion capture and EMG data and ran dynamic simulations with individualized inertial models of the arm to compute muscle torques and their components. We first tested the generalizability of the conclusions from Olesh et al. [23] by comparing principal components obtained from muscle torques with EMG profiles across limbs and workspaces. Here PCA derived scores that represent the projection of each EMG profile in the principal component space. We applied PCA with the assumption of invariant activation profiles to obtain temporal synergies as described above. The scores derived this way will vary across movement directions and muscles but remain constant across time. Then when muscle activity scales together across movement directions we expect the scores of the corresponding muscles to be linearly related. When the muscles whose scores are correlated are agonists, this will capture agonistic co-contraction. Similarly, when the muscles whose scores are correlated are antagonists, this will capture antagonistic co-contraction. Overall, correlations between muscle scores will reveal co-contraction. The more muscles are co-contracting, the lower is the dimensionality of control. The levels of co-contraction were compared between limbs and workspaces. Using this method, we tested the hypothesis that the dimensionality of control space is shaped by biomechanical constraints.

## Materials and Methods

### Data Collection

The experimental protocol was approved by the Institutional Review Board of West Virginia University (Protocol #1311129283). Perspective participants were recruited through fliers distributed around Morgantown, WV. The data collection started on March 12^th^, 2014, and ended on July 25^th^, 2016. Participants provided a written informed consent prior the start of experiments. A second member of the investigative team witnessed the signing of the consent form.

Nine healthy participants (mean age ± SD = 22.78 ± 0.67 years, 3 females and 6 males, sex assigned at birth) performed a modified center-out reaching task by pointing to visual targets in virtual reality (software Vizard by Worldviz, Oculus Rift) [23]. All participants reported they were right-handed and had no neurological or musculoskeletal conditions that could alter movement. Participants repeated reaching movements between a central target and one of 14 targets located along a sphere, with 8 targets placed equidistantly on a horizontal circle parallel with the floor and 8 targets placed equidistantly on a vertical circle parallel to the body sagittal plane, with 2 targets sharing locations in both circles as in Olesh *et al.* [23]. Participants were instructed to point to targets with the index finger, while holding their hand palm down, as quickly and accurately as they can without moving their trunk and wrist. The distances toward the targets were normalized for each subject based on their arm segment lengths to minimize the inter-subject variability in angular kinematics. Each movement to and from each target was repeated 15 times in a randomized order.

The center-out and return movements were repeated with each arm starting at one of two locations of the central target. The central target was placed at the same distance relative to the participant’s shoulder scaled to their arm length. This further minimized the inter-subject variability in angular kinematics. In the lateral starting position, the central target was placed at a location that positioned the shoulder at 0 angle of all degrees of freedom and elbow at 90 degrees, so that the upper arm was parallel to the trunk and the forearm was parallel to the floor (Fig 2, pictograms in top corners, view from above). In the medial starting position, the central target was placed at a location that positioned the hand at the midline of the body and at the same distance from the trunk as in the lateral starting position (Fig 2, pictograms in top center, view from above). The medial starting position was the same for reaches with both right and left arms representing common medial workspace. Thus, four conditions were created based on the location of the central target and the arm used to make reaches, i.e., LLat and RLat for reaches by left and right arm respectively in their respective lateral workspaces and LMed and RMed for the reaches by left and right arm respectively in the common medial workspace.

**Figure 2.**
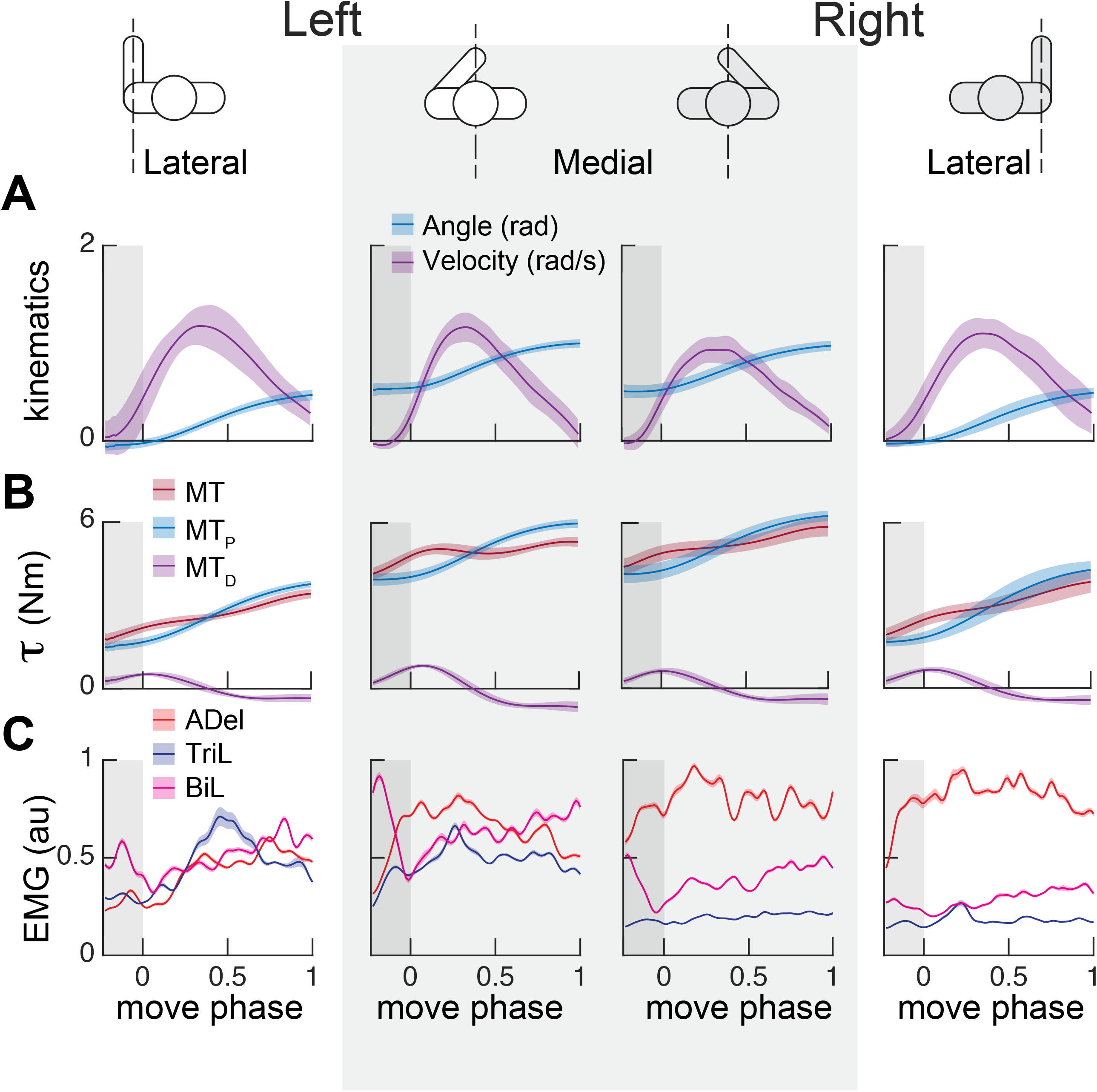
Examples of kinematics, dynamics, and muscle activity profiles during reaching. Each plot shows profiles per condition averaged across repetitions (n = 15) of the same reaching movement in one direction by one participant. (**A**) Profiles (solid lines) and standard deviation (shaded areas) of shoulder flexion-extension angle and angular velocity. (**B**) Shoulder flexion-extension torque (t) profiles that caused the movements in (**A**). (**C**) Normalized electromyography (EMG) profiles of anterior deltoid (ADel), triceps long (TriL), and biceps long (BiL) that accompanied the movements in (**A**). Shaded areas show the standard error for the mean across repetitions (n = 15) of the same movement.

Movement was recorded and visualized in virtual reality using the Impulse system (PhaseSpace Inc) at the temporal resolution of 480 Hz. We tracked the location of 9 light emitting diodes placed on the bony landmarks of the trunk, arm, and hand (Fig 1B) with 8 cameras as described in detail in Olesh et al. [23]. EMG was recorded at the temporal resolution of 2,000 Hz using MA400-28 (MotionLab Systems). EMG was captured from 12 arm muscles: the clavicular head of pectoralis (Pec), teres major (TerM), anterior deltoid (ADel), posterior deltoid (PDel), the long and lateral heads of triceps (TriL and TriS), the short and long heads of biceps (BiS and BiL), brachioradialis (Brd), flexor capri radialis (FCR), flexor carpi ulnaris (FCU), and extensor capri radialis (ECR). EMG recordings were temporally synchronized with the motion capture and virtual reality systems during reaching as described in Talkington *et al.* [28].

### Data Analysis

All data analysis was carried out in MATLAB 2023b (MathWorks Inc). Motion capture data were low pass filtered at 10 Hz and interpolated with a cubic spline. Arm kinematics was obtained from motion capture by defining local coordinate systems representing the trunk, arm, forearm, and hand. Joint angles representing 5 DOFs of the arm were derived using linear algebra, namely shoulder flexion-extension, shoulder abduction-adduction, shoulder internal-external rotation, elbow flexion-extension, forearm pronation-supination, and wrist flexion-extension (Fig 2A, example for one DOF). Active muscle torques and their components were calculated from kinematic data using a dynamic model of the arm with 5 DOFs (Simulink, MathWorks Inc) as described in [23]. The individual height, weight, and segment lengths together with the published anthropometric proportions were used to scale the model inertia to individual body sizes [29]. Inverse simulations were run with the angular kinematics as input and the active torques produced by muscles, termed muscle torques (MT), as output. Simulations were ran with and without gravity force to estimate the components of muscle torque responsible for supporting the arm against gravity (postural component of muscle torque or MT_P_) and for intersegmental dynamics compensation (dynamic component of muscle torque or MT_D_) as described in detail in Olesh *et al.* [23]. The simulations were run for each trial. Torque profiles were normalized in time, averaged per movement direction per DOF and per subject, and down sampled to 100 samples (Fig 2B, example for one DOF). Maximum torque amplitudes were determined per DOF per participant’s arm and used to scale the amplitudes of averaged profiles across movement directions and workspaces.

EMG recordings were high-pass filtered at 20 Hz to remove motion artifacts, rectified, and low-pass filtered at 10 Hz [30]. Next, EMG recordings were normalized to movement duration, down sampled to 100 samples, and averaged per movement direction. Maximum contraction values were determined for each muscle and participant across averaged EMG timeseries per movement direction and used to scale the amplitudes of averaged profiles (Fig 2C).

Principal Component Analysis (PCA) was used to reduce the dimensionality of EMG and torque data similar to Olesh *et al.* [23]. Normalized and demeaned EMG profiles from 12 muscles of 28 center-out reaching movements (14 center-out and 14 return movements between each pair of targets) were combined into one matrix, ***A_n×m_***, where n = 100 (time) and m = 336 (movement directions x muscle). PCA was applied to the data from each of the four conditions (LLat, RLat, LMed, and RMed) separately. Thus, four ***A_n×m_*** matrices were obtained for each subject. Similarly, normalized and demeaned muscle torque profiles (MT, MT_P_, and MT_D_) from 5 DOFs of 28 center-out reaching movements were combined into one matrix, ***B_n×m_***, where n = 100 (time) and m = 140 (movement directions x DOF). Since the analysis included three types of torques (MT, MT_P,_ MT_D_) in the four conditions, twelve ***B_n×m_*** matrices were obtained for each subject. PCA was performed using *pca* function from MATLAB Statistics and Machine Learning Toolbox. Each matrix was centered, and a default singular value decomposition algorithm was applied to obtain principal component eigenvectors (Fig 3A, B), the variance accounted for (VAF) by each principal component (Fig 3C), and scores that are the representations of the matrix in the principal component space. It is important to note that reaching in the same direction leftward or rightward from the starting location rely on different muscles due to the bilateral symmetry of our body. That is why the reaches that include leftward movements to the targets on the left from the central target performed by the left hand were compared to the mirror rightward movements to the right from the central target performed by the right hand and vice versa.

**Figure 3.**
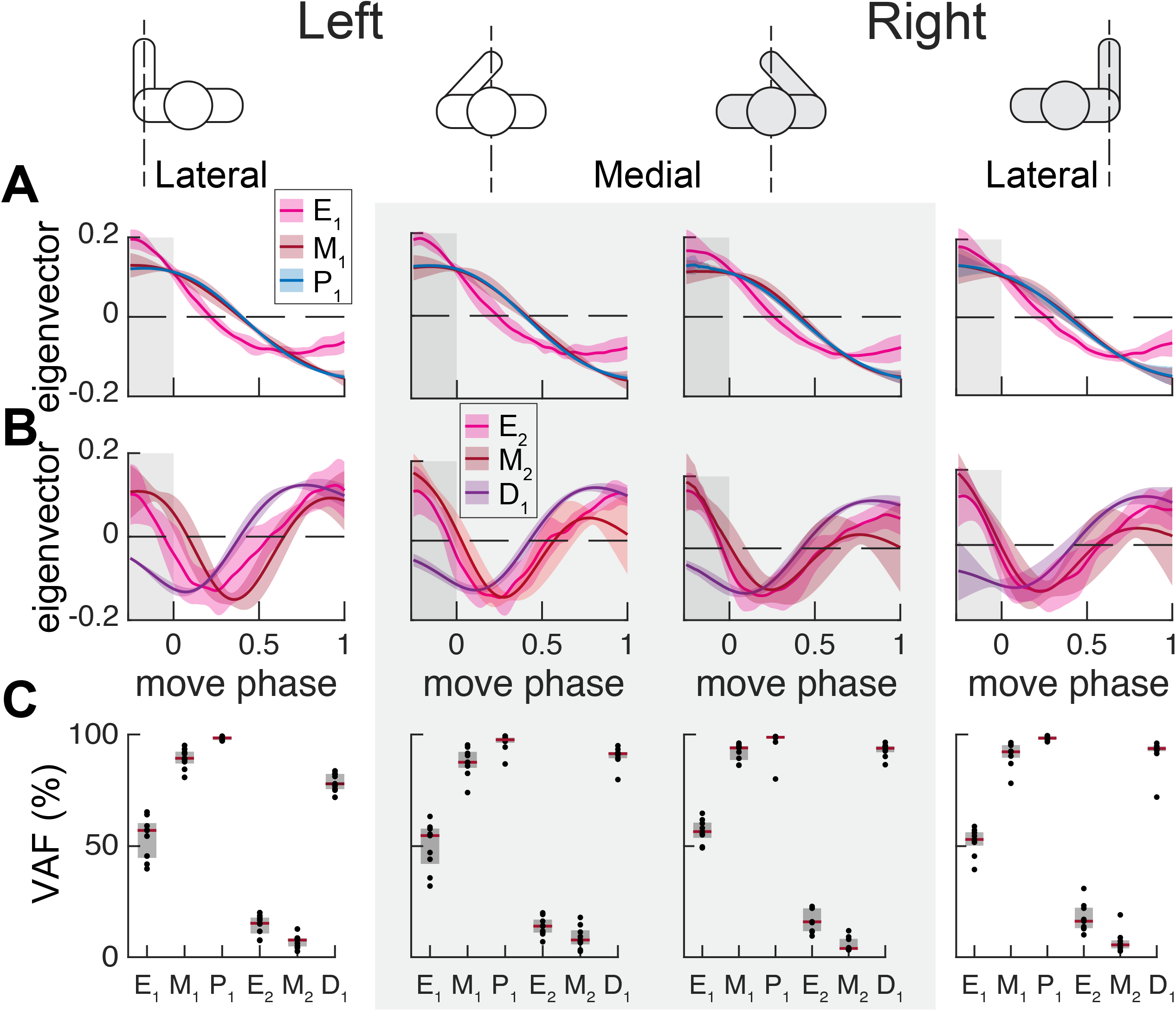
Principal components of EMG and muscle torques. Data from the four conditions indicated by the pictograms are shown in plots arranged in columns. The data is from 9 participants who performed all 4 conditions. (**A**) Solid lines show profiles of the 1^st^ principal component obtained from EMG (E_1_), muscle torque (M_1_), and the postural component of muscle torque (P_1_). (**B**) Solid lines show profiles of the 2^nd^ principal component obtained from EMG (E_2_) and muscle torque (M_2_), and the 1^st^ principal component obtained from the dynamic component of muscle torque (D_1_). (C) Variance accounted for (VAF) by principal components obtained from EMG and muscle torques.

### Statistics and Reproducibility

The first test of the generalizability of conclusions compared the profiles of the 1^st^ and 2^nd^ principal components obtained from EMG to the principal components obtained from muscle torques using a correlation analysis (*corr* function in MATLAB Signal Processing Toolbox). We used repeated measures analysis of variance (RM ANOVA, *ranova* function in MATLAB Statistics Toolbox) to test the null hypotheses that the coefficients of determination (R^2^) from all conditions and data types come from the same distribution. The between-subject factor was Sex (assigned at birth), the within-subject factors were Conditions (4 levels at LLat, RLat, LMed, and RMed) and Components (4 levels comparing E_1_ vs. M_1_, E_2_ vs. M_2_, E_1_ vs. P_1_, and E_2_ vs. D_1_). Post-hoc multiple comparisons were done using *multcompare* function in MATLAB Statistics Toolbox. We applied Bonferroni-Sidak correction to correct the alpha for familywise error, so that the adjusted alpha for the comparison between 6 combinations of 4 conditions = 0.0085 and for the comparison between 6 meaningful combinations of two principal components = 0.0085. The ethnicity – based statistical analysis was not performed due to small sample size.

To test the hypothesis, we evaluated how muscles co-activate across changing starting postures and movement directions. The rationale is that reaching in different directions is caused by different amplitudes of postural or dynamic torques. Therefore, given that the EMG variance is captured well by these torques (1^st^ test above), the corresponding features in EMG are also changing with the amplitudes of the torques. For example, if the amplitude of shoulder flexion postural torque was larger during movement upward compared to the movement forward, then the 1^st^ principal component score of the anterior deltoid (AD, shoulder flexor) EMG will also be larger during movement upward compared to the movement forward. If the scores of multiple muscles change together across movement directions, then these muscles are co-contracting together and can be activated by a single control signal. The advantage of this method of calculating co-contraction is that it does not depend on the assumptions underlying the scaling of EMG amplitude. As long as the relative changes in the EMG amplitudes across movement directions are preserved, the method works.

This method to calculate co-contraction was implemented using a linear correlation analysis (*corr* and *corrplot* functions in MATLAB Statistics and Econometrics Toolboxes) aimed at quantifying the linear relationships between principal component scores from pairs of muscles across movements. First, we extracted the scores of the 1^st^ principal component from EMG and concatenated them into matrix ***C_n×m_***, where n = 28 (movement directions) and m = 12 (muscles) per condition per subject. Positive correlations indicate co-contraction of muscles, which negative correlations indicate reciprocal activation of muscles, i.e., if the EMG activity related to postural or dynamic torques increases in one muscle it decreases in another muscle. We evaluated the statistical significance of the linear relationships represented by the Coefficients of Determination (R^2^) between scores of 66 muscle pairs (Fig 5). We applied Bonferroni-Sidak correction to correct the alpha for familywise error, so that the adjusted alpha = 0.0008. We repeated this analysis for the 2^nd^ principal component from EMG that captures the dynamic torque feature in EMG.

**Figure 4.**
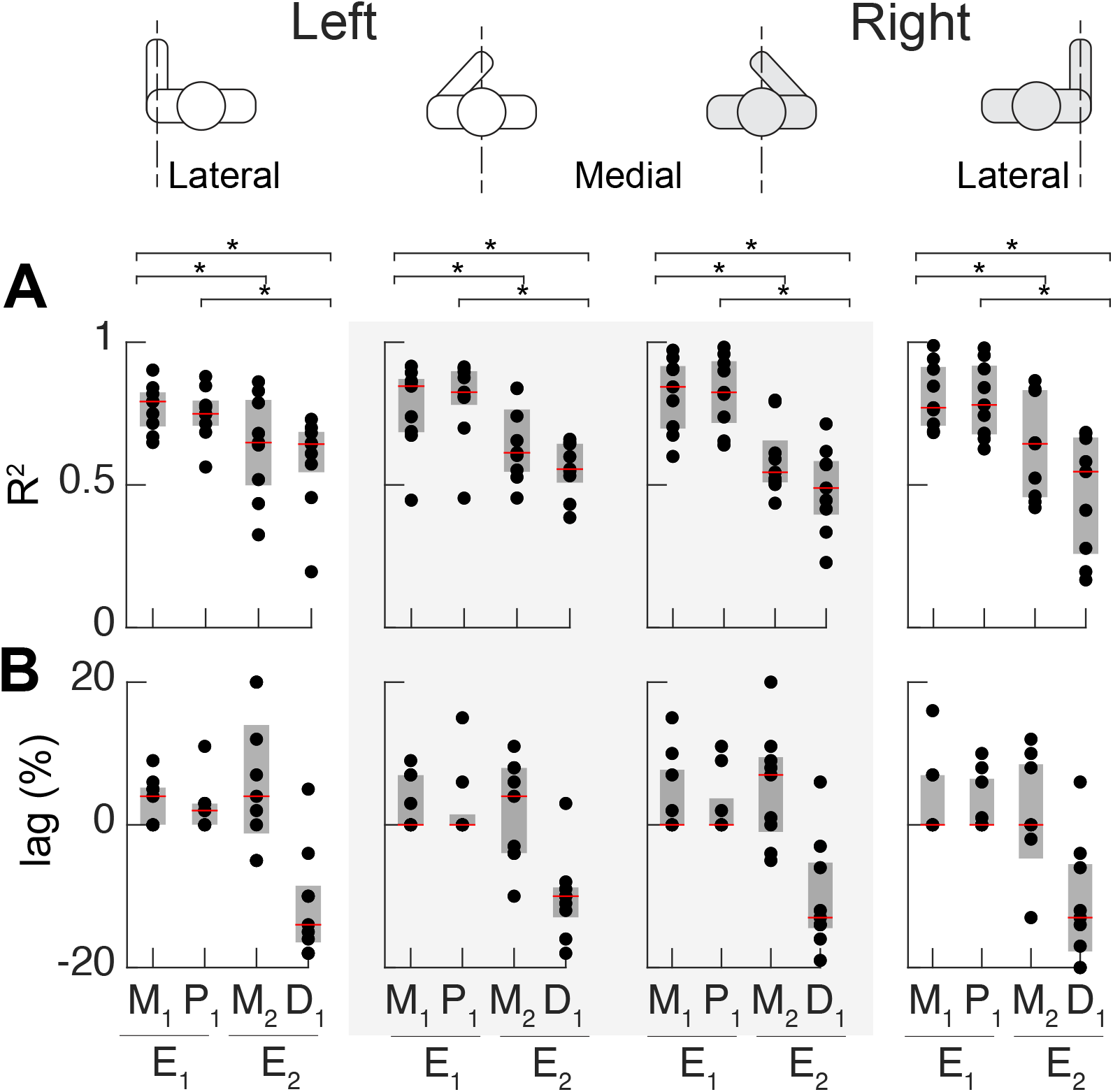
Correlations between the temporal profiles of principal components derived from EMG and muscle torques. Data (N = 9) from the four conditions indicated by the pictograms are shown in plots arranged in columns. (A) Dots show individual R^2^ values from cross-correlations between profiles, red lines show median values, and grey boxes show interquartile ranges. Brackets with stars indicate significant relationships with correction for familywise error. (B) Dots show individual lag values from cross correlations, red lines show median values, and grey boxes show interquartile ranges. Open dots are outliers. E_1_ and E_2_ are the 1^st^ and 2^nd^ principal components respectively obtained from EMG; M_1_ and M_2_ are the 1^st^ and 2^nd^ principal components respectively obtained from muscle torques; P_1_ and D_1_ are the 1^st^ principal components obtained from the postural and dynamic components of muscle torque respectively.

**Figure 5.**
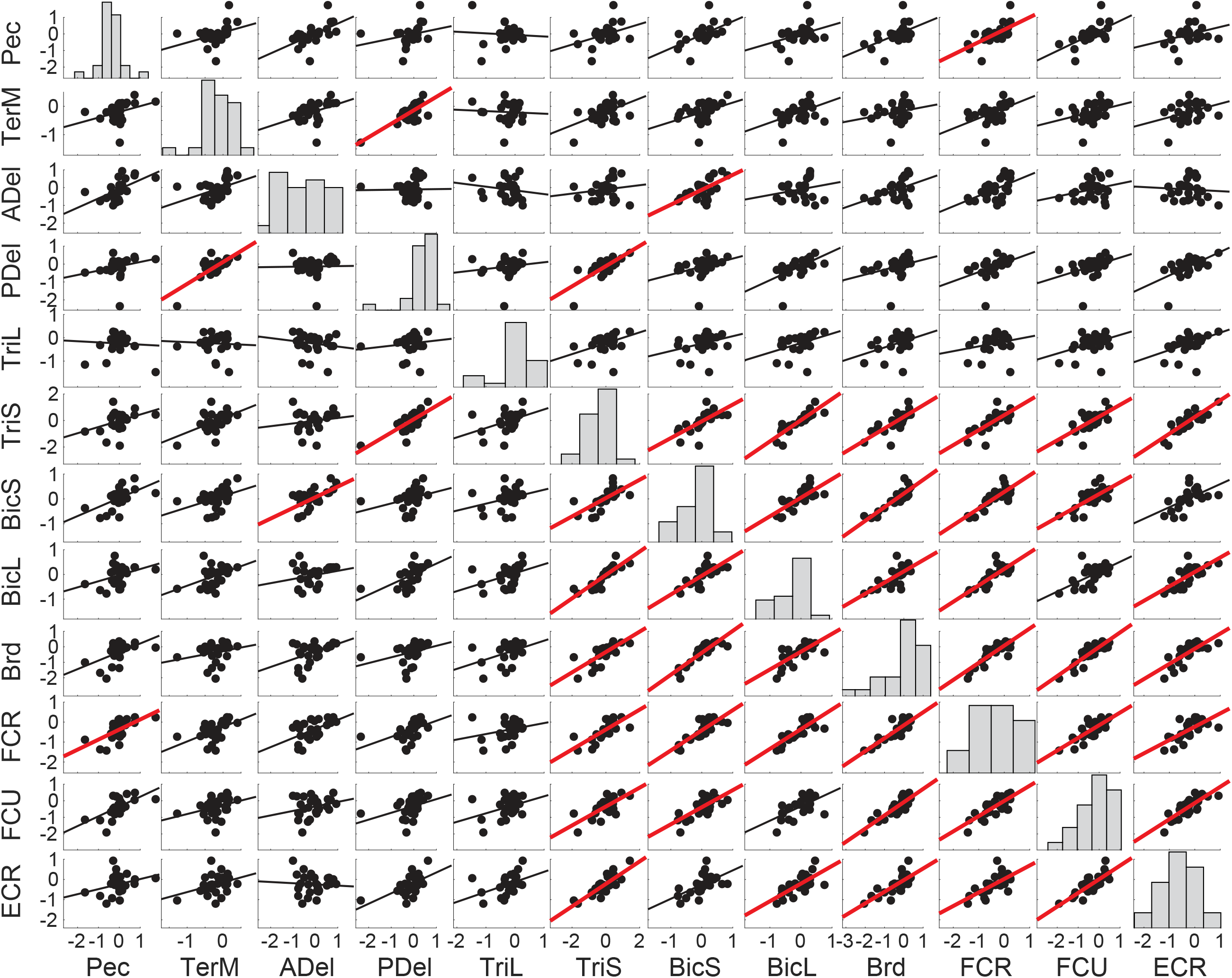
Example linear regressions between scores across reaching directions. Example regression matrix between E_1_ scores of 12 muscles across 28 right-handed reaches within the medial workspace (RMed condition) for a single participant. The coordinates of each dot represent principal component scores for two muscles for a single movement direction. Histograms along the diagonal show the distribution of the scores for a given muscle across reaching directions. Solid lines show least-squares linear regression, red lines indicate significant relationships with correction for familywise error, so that the adjusted alpha = 0.0008.

## Results

Participants performed center-out and return movements with each arm starting at one of two locations of the central target. Four conditions were created based on the location of the central target and the arm used to make reaches, i.e., LLat and RLat for reaches by left and right arm respectively in their respective lateral workspaces and LMed and RMed for the reaches by left and right arm respectively in the common medial workspace. All participants were able to complete the center out task with either left or right arms with consistent kinematics (Fig 2A). Due to bilateral symmetry, we expect that movements with left and right arms with the same starting positions and kinematics in forward, backward, up, or down directions rely on similar forces produced by the same muscles. For the same reason movements leftward or rightward from the starting positions by one limb will be similar to the rightward and leftward movements respectively by the other limb. The forces underlying these movements were inferred from motion capture using dynamic modeling that calculated muscle torques about each DOF of the shoulder, elbow, and wrist joints. These torques were further subdivided into a linear combination of postural and dynamic torques. The muscle torque profiles accompanying the corresponding movements had very similar trajectories across conditions (Fig 2B). However, there were static offsets in the postural torques underlying matching movement directions. For example, holding the initial position at the beginning of the movement required larger muscle torques about the right shoulder flexion/extension DOF (RMed and RLat conditions) compared to that for the left shoulder in matching conditions (LMed and LLat conditions respectively, Fig 2B). This is likely due to the motor redundancy or the multiple combinations of joint angles and torques that can result in the same position of the hand.

From our earlier work we knew that important features of muscle activity can be captured by the postural and dynamic torques [23]. However, in that study we focused our analysis on the movements performed by the dominant arm in the lateral workspace. Here the analysis encompasses different workspaces and both arms, which enabled us to evaluate the generalizability of this conclusion. We found that intra- and inter-subject variability of EMG profiles were very low. The average standard deviation of EMG profiles across movement repetitions for muscles in the dominant right arm (normalized to maximal EMG of each muscle) was 12% ± 1% (standard deviation across subjects, SD). The corresponding variability of EMG of muscles in the non-dominant left arm was 22% ± 2%. This shows that the muscle activation profiles were more consistent across repetitions of the same movement type for the dominant arm compared to the non-dominant arm. Moreover, the EMG profiles also broadly reflected the differences in the joint torques across conditions (Fig 2C). In the example for the reach forwards and up in Fig 2, the amplitude of anterior deltoid activity increased in parallel with the increases in the postural component of the shoulder flexion muscle torque across conditions (Fig 2, MT_P_ and ADel). Furthermore, the initial propulsive flexion moment in the dynamic muscle torque profile was preceded by a burst in the activation of the long head of biceps (Fig 2, MT_D_ and BiL, end of the burst is visualized). However, EMG profiles are notoriously noisy and difficult to interpret. Therefore, in the following analysis we will broadly test the generalizability of conclusions from Olesh et al. [23] by focusing on the salient features in EMG obtained with principal component analysis (PCA; see details in Methods).

The temporal profile of the 1^st^ principal component (eigenvector E_1_) obtained from EMG matched the 1^st^ principal component (M_1_) obtained from muscle torques (MT) and the 1^st^ principal component (P_1_) obtained from the postural torques only (MT_P_; Fig 3A). The 1^st^ principal component captures the gradually increasing or decreasing EMG or torque profiles associated with the changes in posture from the starting position to the final position at the reach target. The high variance captured by P_1_ indicates that the profiles of muscle torques were very similar across joints and DOFs, indicating a high degree of intralimb coordination. As expected, the variance accounted for (VAF) by the 1^st^ principal component in EMG (E_1_ = 53%) was lower than that for muscle torque (M_1_ = 90%; Fig 3C) due to higher variability of EMG profiles compared to torque profiles.

The temporal profile of the 2^nd^ principal component (E_2_) obtained from EMG matched the 2^nd^ principal component (M_2_) obtained from muscle torques and the 1^st^ principal component (D_1_) obtained from the dynamic torques only (MT_D_) although with a phase shift (Fig 3B). This feature captures the acceleration/deceleration-related changes in torques that underly bursts in EMG profiles. The VAF by the 1^st^ principal components obtained from muscle torques (M_1_, P_1_, and D_1_) was very high (Fig 3C), the mean values across conditions and all participants were 90% for M_1_, 97% for P_1_, and 88% for D_1_ (Table 1 contains values per condition).

**Table 1.**
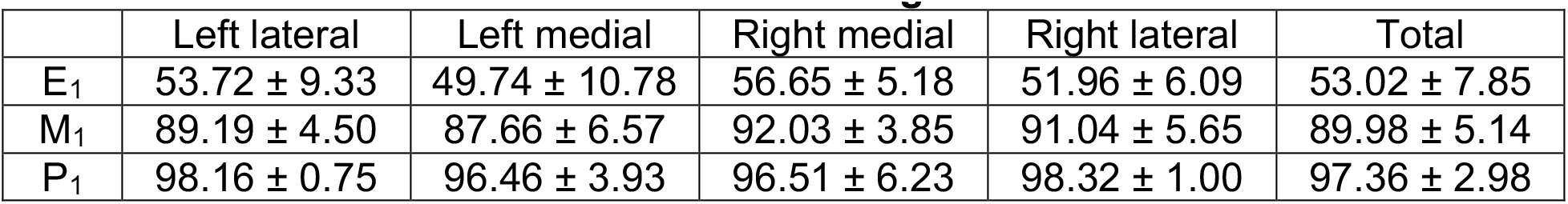
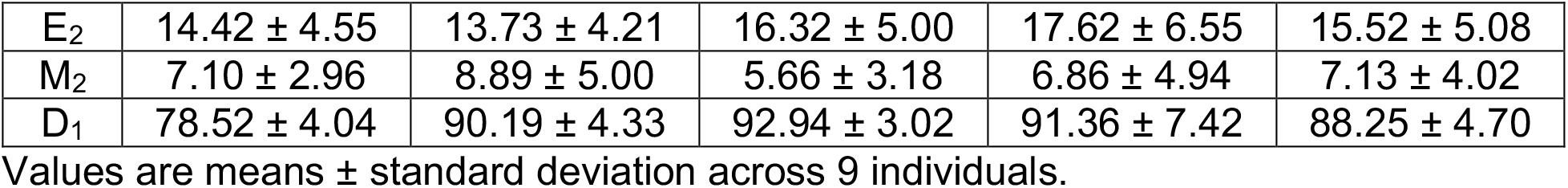
Variance accounted for across reaching conditions.

Interestingly, P_1_ captured more variance across conditions and DOFs than M_1,_ similarly D_1_ captured more variance across conditions and DOFs than M_2_ despite similar profiles. This shows that subdividing muscle torques into postural and dynamic components results in more reliable features compared to those obtained with PCA of muscle torques. More VAF in EMG was captured by the 2^nd^ principal component (E_2_ = 16%) than that in muscle torque (M_2_ = 7%; Fig 3C). This indicates that highly stereotypical muscle torques during reaching movements explain a high portion of the overall variance in EMG profiles. The same conclusion was reached in Olesh et al. [23] from a smaller subset of data. Moreover, our results further support the idea that E_1_ captures the muscle recruitment necessary for generating forces to counteract gravity, while E_2_ captures the muscle recruitment necessary for generating forces for accelerating and decelerating the arm toward the target. This generalizes our earlier conclusions that the static and phasic components of EMG envelope from a given muscle reflects the muscle’s contribution to gravity compensation and propulsion toward the target respectively.

The degree of similarity between features obtained with PCA across torques and EMG was analyzed using cross correlation analysis as described in Methods. One-tailed repeated measures analysis of variance (RM ANOVA) on the coefficients of determination (R^2^) between principal component profiles showed a significant main effect (F(15, 105) = 9.36, *p* = 0.00) but accounting for sex reduced differences between conditions and components (F(15, 105) = 1.21, p = 0.27). Two-tailed post-hoc tests have shown no differences between males and females and between conditions (Table 2). There were significant differences between signal types from which components were derived indicating that the temporal profiles of E_1_, M_1_, and P_1_ were more similar to each other than to E_2_, M_2_, or D_1_ and vice versa (Table 3; Fig 4A). This further supports the idea that the 1^st^ and 2^nd^ principal components of EMG represent the contribution of the muscles to postural and dynamic torques. These conclusions generalize across both limbs and across medial and lateral workspaces. The profiles of postural and dynamic torques captured most of variability in muscle activity profiles. Thus, this method of EMG decomposition can be used to remove noise from EMG signals and increase the interpretability of surface EMG recordings.

**Table 2.** Post-hoc differences between the coefficients of determination across males and females and across limbs and workspaces across 9 participants.

| Sex |  | MSE ± SE | p value |
| --- | --- | --- | --- |
| F | M | -0.10 ± 0.04 | 0.06 |
| Conditions |  | MSE ± SE | p value |
| LLat | LMed | -0.02 ± 0.03 | 0.95 |
| LLat | RLat | 0.01 ± 0.03 | 0.96 |
| LLat | RMed | 0.01 ± 0.04 | 1.00 |
| LMed | RLat | 0.03 ± 0.04 | 0.88 |
| LMed | RMed | 0.02 ± 0.04 | 0.91 |
| RLat | RMed | -0.01 ± 0.02 | 0.98 |

MSE – mean squared error, SE – standard error of the mean. Adjusted alpha = 0.0085.

**Table 3.** Post-hoc differences between the coefficients of determination based on EMG- and torque-based components across 9 participants.

| Components |  | MSE ± SE | p value |
| --- | --- | --- | --- |
| E<sub>1</sub>/M<sub>1</sub> | E<sub>1</sub>/P<sub>1</sub> | -0.01 ± 0.01 | 0.9036 |
| E<sub>1</sub>/M<sub>1</sub> | E<sub>2</sub>/M<sub>2</sub> | 0.19 ± 0.04 | **0.0048** |
| E<sub>1</sub>/M<sub>1</sub> | E<sub>2</sub>/D<sub>1</sub> | 0.29 ± 0.05 | **0.0040** |
| E<sub>1</sub>/P<sub>1</sub> | E<sub>2</sub>/M<sub>2</sub> | 0.20 ± 0.05 | 0.0155 |
| E<sub>1</sub>/P<sub>1</sub> | E<sub>2</sub>/D<sub>1</sub> | 0.30 ± 0.06 | **0.0066** |
| E<sub>2</sub>/M<sub>2</sub> | E<sub>2</sub>/D<sub>1</sub> | -0.10 ± 0.05 | 0.2367 |

MSE – mean squared error, SE – standard error of the mean. Bold p values show significant difference with adjusted alpha = 0.0085.

We have observed a consistent phase lead of the D_1_ components onto the E_2_ component (Fig 4B). This phase lead can potentially be taken advantage of to predict the changes in the underlying muscle forces or contractions from dynamical simulations prior to observable motion and improve performance of real-time control applications.

To test the hypothesis that the dimensionality of control space is shaped by biomechanical constraints we quantified co-contraction using correlations between PCA scores from EMG. The method relies on the established relationships between torque and EMG components. These relationships imply that for movements requiring a larger change in the postural torques, higher scores for the principal component representing that feature (E_1_) will be observed in one or more muscles in the same movement. If the scores change together across movement directions in several muscles, these muscles are co-contracting and can be activated by the same neural control signal. This means that if the primary features in EMG (E_1_ and E_2_) represent low dimensional control signals, we expect significant linear correlations between corresponding scores of multiple muscles. We performed this linear correlation analysis per arm and workspace (Fig 5 shows an example for one condition and participant).

The scores represent the projection of EMG onto the principal component vectors. When the scores are close to 0, this means that the components do not capture much variance in the corresponding EMG profile. For all combinations of muscles, movements, and participants, about 21% of the E_1_ scores were low, i.e., values were below 5% of maximal score (RLat: 24%, LLat: 19%, LMed: 17%, and RMed 22%). About 26% of E_2_ scores were low (RLat: 27%, LLat: 26%, LMed: 22%, and RMed: 28%). The low score means that the EMG profile for a given muscle in a given movement is not captured well by PCA. Therefore, no relationships between the low score values are expected, as seen for example in the insignificant correlation between low scores of BicL and ADel in the example shown in Fig 5. However, we did observe multiple significant correlations between E_1_ scores in certain conditions. For reaching within the left lateral workspace, 61% of muscles (40 muscle pairs out of 66) in the non-dominant arm showed moderate correlations between their E_1_ scores (R^2^ ≥ 0.5; Fig 6A). For reaching within the medial workspace, 41% of muscles (27/66 pairs) in the non-dominant arm showed moderate correlations between their E_1_ scores (Fig 6B). In contrast, only 11% of muscles in the dominant arm showed moderate correlations when reaching within the right lateral workspace (Fig 6C). For reaching within the shared medial workspace, 29% of muscles in the dominant arm showed moderate correlations (Fig 6D), closer but lower than that for the non-dominant arm (Fig 6B). Overall, more correlations were observed between E_1_ scores in muscles of the non-dominant arm in 7 out of 9 individuals (Figs S1-S9 top row). This shows that the forces needed to support the arm against gravity can be generated by few control signals evidenced by the large number of co-contracting muscles, particularly in the non-dominant arm. In contrast, there were much fewer muscle pairs with consistently correlated E_2_ scores (Fig 7). This was true for all individuals (Figs S1-S9 bottom row). This suggests that the generation of the propulsive forces requires higher dimensionality of control than that of the postural forces evidenced by the lack of consistent co-contraction patterns across muscles. Overall, these results support the hypothesis that the biomechanics shapes the dimensionality of control space.

**Figure 6.**
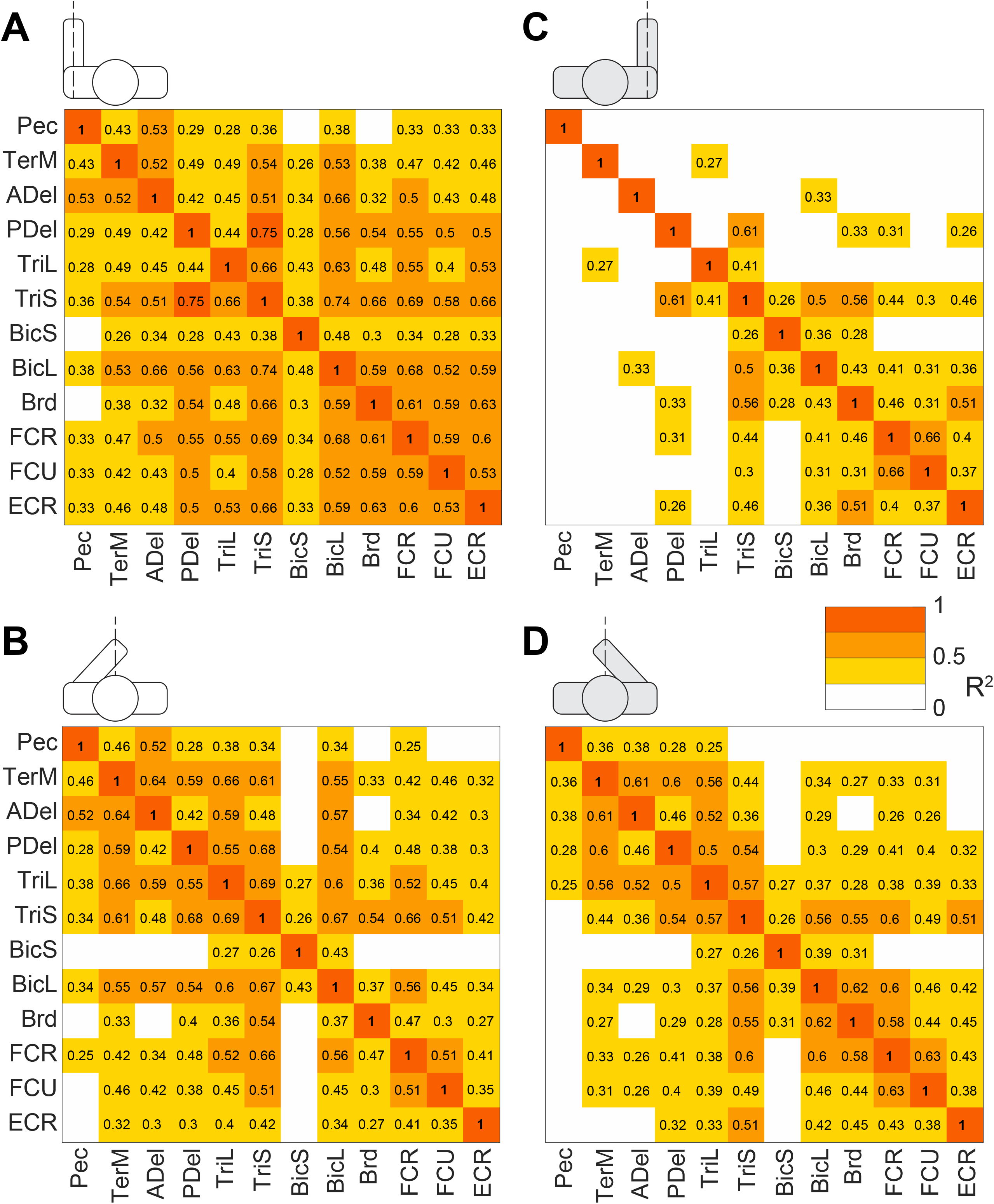
Gravity-related muscle co-contraction across all conditions. Heatmaps show coefficients of determination (R^2^) from regressions shown in Fig 4 averaged across 9 participants. Red and orange colors represent strong and moderate relationships respectively between E_1_ coefficients across movement directions. Pictograms indicate conditions, i.e., LLat (A), LMed (B), RLat (C), and RMed (D). Muscles are abbreviated as described in Methods.

**Figure 7.**
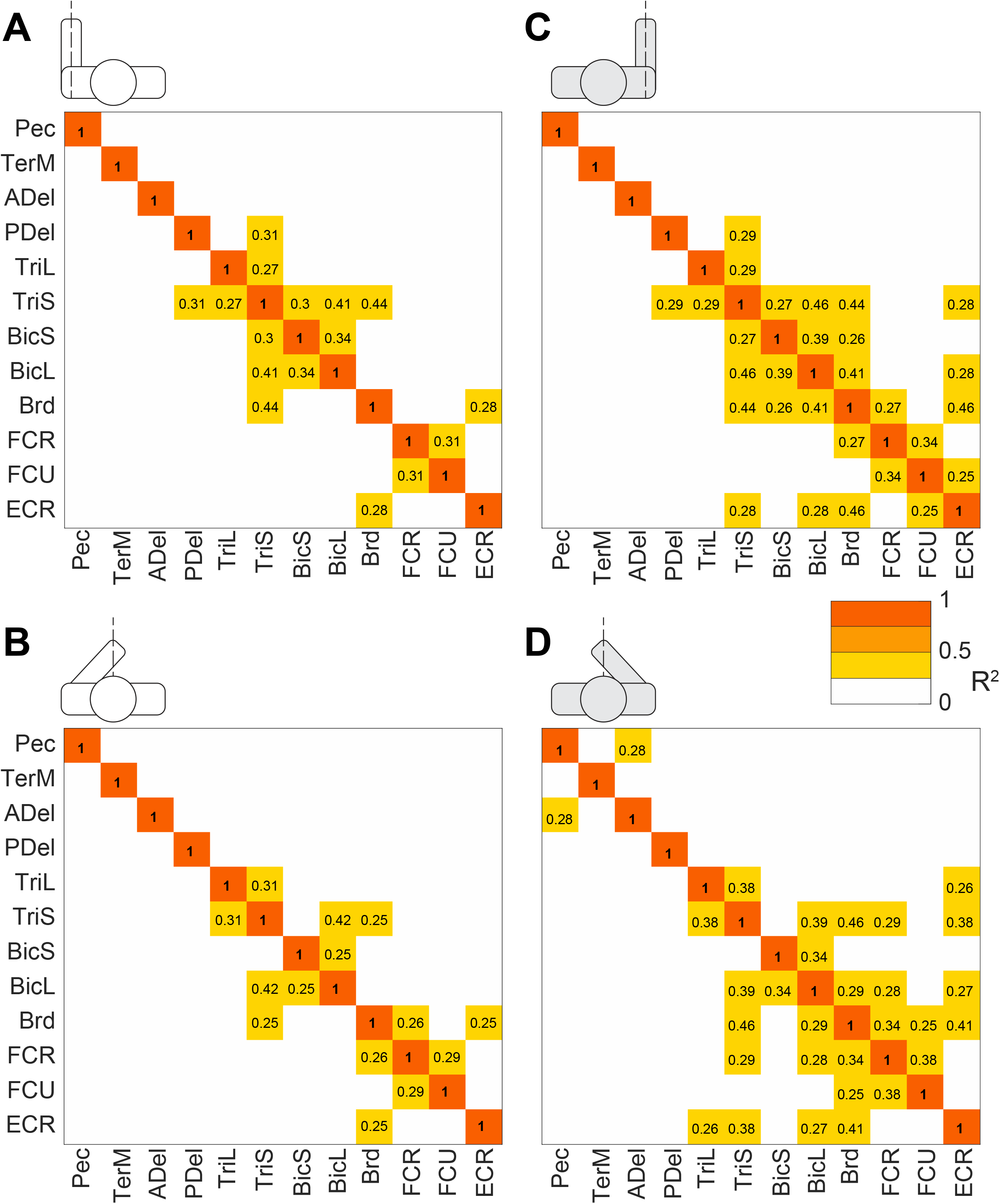
Dynamic torque-related muscle co-contraction across all conditions. Heatmaps show coefficients of determination (R^2^) from regressions shown in Fig 4 averaged across 9 participants. Red, orange and darker blue colors represent moderate and strong relationships between E_2_ coefficients across movement directions. Pictograms indicate conditions, i.e., LLat (A), LMed (B), RLat (C), and RMed (D). Muscles are abbreviated as described in Methods.

Correlations between scores were overwhelmingly positive, indicating that not only agonistic but also antagonistic muscles, such as biceps and triceps, changed their activity together. Note that in these reaching movements with the arm pronated, the muscles performing the antigravity action would be ADel, BiL, Brd, and ECR, while the rest would produce joint moments in the direction of gravity (Fig. 1B). There were the most correlations between E_1_ scores in the non-dominant arm in the LLat condition. Two salient muscle groups comprised proximal muscles (TerM, ADel, PDel, TriL, and TriS) and distal muscles (BiL, Brd, FCR, FCU, and ECR), both highly coupled (Fig 6A). In contrast, the correlations between E_1_ scores in the dominant arm in the corresponding RLat condition consisted of three much smaller muscle groups (PDel/TriS//BicL/Brd, Brd/ECR, and FCR/FCU) with only the first two groups coupled through Brd (Fig 6C). The three small clusters were the only consistent groups of coactivating muscles across both workspaces and both limbs. This shows that both agonistic and antagonistic muscles are recruited to produce forces that support the limb against gravity, indicating that limb stiffness can help reduce the dimensionality of neural control space.

The co-contraction patterns for generating dynamic torques were much less consistent across conditions and individuals. The correlations between E_2_ scores in the dominant arm in the RLat condition, only 4 pairs of muscles were consistently positively correlated across individuals with <0.5 R^2^ (R^2^ for BicL/TriS = 0.46, Brd/TriS = 0.44, Brd/BicL = 0.45, and ECR/Brd = 0.46). In individuals the patterns of co-contraction were even less consistent (Figs S1-S9, bottom row). There were 6 significantly correlated muscle pairs in RLat condition that were similar across at least 5/9 subjects (Brd/ECR, FCR/FCU, Brd/BicL, Brd/TriS, BicL/BicS, and BicL/TriS) and 3 significantly correlated muscle pairs in RMed condition (Brd/ECR, Brd/TriS, and BicL/TriS). For the non-dominant arm, there were 2 significantly correlated muscle pairs in LLat condition that were similar across 5/9 subjects (Brd/TriS and BicL/TriS) and 1 significantly correlated muscle pair in LMed condition (BicL/TriS). The co-activation of most of these muscle groups, with the exception of Brd/TriS and BicL/TriS, was agonistic. Moreover, some instances of negative correlations between antagonistic muscle pairs were observed in individuals (ADel/TerM; PDel/Pec, BicL/PDel, and ECR/FCU; Figs S1-S9 bottom row). This shows that more complex control signals coordinating both co-contraction and reciprocal activation of muscles are needed to produce forces that propel the limb in different directions from different starting locations.

## Discussion

We observed that highly stereotypical muscle torques during reaching movements explain a high portion of the overall variance in EMG profiles in multiple arm muscles. We have shown that the first principal component of EMG captures the muscle recruitment necessary for generating forces to counteract the force of gravity, while the second principal component of EMG captures the muscle recruitment necessary for generating dynamic forces for accelerating and decelerating the arm toward the target (Figs 3 and 4). These conclusions, originally derived from a smaller subset of movements by the dominant arm in one area of reaching workspace [23], are generalizable to movements by the non-dominant arm and across a larger reaching workspace that overlaps between both arm arms. This is useful for understanding the dimensionality of neural control space.

We have also observed correlations between scores of muscle pairs across movements indicating co-contraction but which and how many muscles co-contracted varied under deferent conditions (Figs 6, 7, and S1-S9). We have shown that the forces needed to support the arm against gravity can be generated by few control signals evidenced by the substantial number of co-contracting muscle pairs, particularly in the non-dominant arm. This is due to the force of gravity always pointing in the same direction, so that the counteracting forces can be produced by consistent groups of muscles. We have also shown that the generation of the propulsive forces require higher dimensionality of control than that of the postural forces evidenced by the lack of consistent co-contraction patterns across muscles. This is likely due to the direction-specific forces needed to make the movements to the different targets, which require different combinations of muscle forces. These results support the second hypothesis that the biomechanics shapes the dimensionality of control space.

It is possible that the reason we have not obtained consistent co-contraction patterns across movements is that the surface EMG was too noisy and variable across the different movements and individuals. However, our variability data reported in Results shows that the temporal profiles of EMG were very consistent across all these conditions, especially for the dominant arm with the fewest observed co-contractions. Another evidence of the consistency of EMG profiles is the large percentage of variance that is captured in EMG by the first two principal components (Fig 3) and by the components obtained from muscle torques (Fig 4), which are highly stereotypical during reaching movements. Therefore, it seems highly unlikely that the inter-movement or inter-subject variability can explain the lack of consistent co-contraction across different conditions.

The neural control strategies underlying muscle recruitment during different phases of movement can be quantitatively described through not only muscle torques but also mechanical impedance. Muscles are organized in a complex pattern to generate opposing moments around each joint DoF as determined by intrinsic muscle properties [31] and muscle moment arms. Therefore, neural muscle recruitment causes not only muscle torques but also the counterbalanced forces that define the stiffness and viscosity components of impedance, which in turn determine the reaction of the musculoskeletal system to perturbations [32]. The impedance parameters that are under the control of the nervous system can be measured by assessing changes in the co-contraction of antagonistic muscles [26]. Both stiffness and viscosity components of mechanical impedance define how far the joints move in response to an applied external force [32]. Muscle recruitment defines the functional aspects of joint impedance that can and do change with task demands [33,34]. Here too we observed that the component of EMG that is responsible for supporting the limb against gravity is present in not only antigravity muscles but also in their antagonists and that the amplitude of this component changes together across movement directions indicating co-contraction (Fig 6). Other studies have shown that the amount of muscle co-activation increases in movements with unstable loads compared to those with stable loads [35] and the impedance of the whole limb increases to stabilize movement when learning novel dynamical tasks [33]. The joint stiffness component of impedance has been shown to be higher during unconstrained movements compared to constrained ones indicating, unsurprisingly, that unconstrained movements are inherently less stable [36]. Limb impedance increases when learning to control an unstable object and when movement stability is challenged by a robot [37,38]. Wrist stiffness increases during unstable dynamical loads compared to stable ones [39] and when the accuracy demands of the task are high [40]. Moreover, we observed more co-contraction in the non-dominant limb compared that in the dominant limb, supporting earlier observations that that there is hemispheric specialization in the compensation for limb dynamics. The non-dominant limb motor control has been shown to be ore specialized for impedance control [41–43] Overall, this suggests that the compensation for gravity load on the arm during reaching is accomplished through a low-dimensional control space by controlling limb stiffness. Low-dimensional control of posture through stereotypical postural adjustments and other reflexes has been shown to involve the reticulospinal tract [18–20] (Fig. 1, green).

Our results also support the idea of internal models embedding body dynamics. Internal models are defined as neural networks in the brain that simulate the dynamics of the body and its interaction with the external world [1,3,44–47]. Forward models predict the consequences of a given action, for example, predicting sensory feedback without delay from an efferent copy of a corticospinal motor command [48,49]. Inverse models capture information in the opposite direction [3,50]. They convert a motor plan into the motor execution output of the motor system that converges on the motoneurons in the spinal cord. The motor execution output is thought to be forces in the intrinsic reference frame that are closely related to individual muscle forces or moments [51,52]. The motor plan is thought to be developed in the premotor areas, where neural activity has many kinematic features in the extrinsic head-centered reference frame resolving the kinematic redundancy problem [53–55]. The motor execution output is thought to originate in the primary motor cortex, where neural activity has many dynamic features in the intrinsic reference frame [51,56–58]. The transformations from motor plan into motor execution may potentially resolve the muscle redundancy problem by embedding biomechanical constraints. These constraints may take the form of embedded muscle anatomy relationships, such as known ratios between moments arms of muscles about a given DOF [59]. Here too we have observed that a large amount of variance in the muscle activity profiles are captured by muscle torques. Interestingly, the component of muscle activity that reflects propulsion was not associated with consistent patterns of antagonistic co-contraction across multiple movement directions (Fig 7). Instead, smaller subsets of mainly agonistic co-contraction and reciprocal activation of antagonists was observed(Figs 7, S1-S9). This provides evidence that control solutions with higher dimensionality are needed for generating these forces. Internal models could support high-dimensional control, because the control solutions are thought to be calculated on the fly by the neural networks that embed the equations of motion and biomechanical constraints. These calculations likely involve the higher-order neural structures whose output converges on the corticospinal tract [21,60–62] (Fig. 1, purple).

## Conclusions

Results show that across multiple movement directions and starting positions and in both arms muscle activity broadly reflects the temporal evolution of muscle torques. We suggest that the muscles generate forces to counteract gravity through a simplified set of control signals, leading to coordinated contraction among various muscles, which adjusts the stiffness of the limb. Conversely, the forces that drive the limb towards a target demand more intricate control signals that vary with the direction of movement.

## Acknowledgements

We would like to acknowledge the contributions of Dr. E.V. Olesh and Dr. A.B. Thomas to the collection and preliminary analysis of the reported data. V.G. was supported by NIGMS grants P20GM109098 and P30GM103503. ASK was supported by a fellowship from NIGMS T32 AG052375. This work was supported in part by the Office of the Assistant Secretary of Defense for Health Affairs through the Restoring Warfighters with Neuromusculoskeletal Injuries Research Program (RESTORE) under Award No. W81XWH-21-1-0138. Opinions, interpretations, conclusions, and recommendations are those of the author and are not necessarily endorsed by the Department of Defense.

## Author Contributions

A.S.K. contributed to conceptualization, data curation, data analysis, and writing. V.G. contributed to conceptualization, methodology development, data curation, data analysis, and writing. All co-authors have read and edited the manuscript and agree with its content.

## Data Availability Statement

The data used for statistical analysis, such as principal components and scores for electromyography and muscle torques, are shared online [63].

## Supporting information

**Supplementary Figures S1-S9. Gravity and dynamic torque-related muscle co-contraction across all conditions for each participant.** Heatmaps show coefficients of determination (R^2^). Red, orange and darker blue colors represent moderate and strong relationships between E_1_ (A, B) and E_2_ (C, D) coefficients across movement directions;. Pictograms indicate conditions, i.e., LLat (A, C) and RLat (B, D). Muscles are abbreviated as follows: the clavicular head of pectoralis (Pec), teres major (TerM), anterior deltoid (ADel), posterior deltoid (PDel), the long and lateral heads of triceps (TriL and TriS), the short and long heads of biceps (BiS and BiL), brachioradialis (Brd), flexor capri radialis (FCR), flexor carpi ulnaris (FCU), and extensor capri radialis (ECR).

